# An improved growth medium for enhanced inoculum production of the plant growth-promoting fungus *Serendipita indica*

**DOI:** 10.1101/857128

**Authors:** Mohamed Osman, Christian Stigloher, Martin J. Mueller, Frank Waller

## Abstract

The plant endophytic fungus *Serendipita indica* colonizes roots of a wide range of plant species and can enhance growth and stress resistance of these plants. Due to its ease of axenic cultivation and its broad host plant range including the model plant Arabidopsis thaliana and numerous crop plants, it is widely used as a model fungus to study beneficial fungus-root interactions. In addition, it was suggested to be utilized for commercial applications, e.g. to enhance yield in barley and other species. To produce inoculum, *S. indica* is mostly cultivated in a complex Hill-Käfer medium (CM medium), however, growth in this medium is slow, and yield of chlamydospores, which are often used for plant root inoculation, is relatively low. We tested and optimized a simple vegetable juice-based medium for an enhanced yield of fungal inoculum. The described vegetable juice (VJ) medium is based on commercially available vegetable juice and is easy to prepare. VJ medium was superior to the currently used CM medium with respect to biomass production in liquid medium and hyphal growth on agar plates. Using solid VJ medium supplemented with sucrose (VJS), a high amount of chlamydospores developed already after 8 days of cultivation, producing significantly more spores than on CM medium. Use of VJ medium is not restricted to *S. indica*, as it also supported growth of two pathogenic fungi often used in plant pathology experiments: the ascomycete *Fusarium graminearum*, the causal agent of Fusarium head blight disease on wheat and barley, and *Verticillium longisporum*, the causal agent of verticillium wilt. The described VJ medium is recommended for a streamlined and efficient production of inoculum for the plant endophytic fungus *Serendipita indica* and might prove superior for the propagation of other fungi for research purposes.

## Background

*Serendipita indica*, first described as *Piriformospora indica* [1], is a facultative root endophytic fungus belonging to the basidiomycotal order *Sebacinales*. Within this clade, life styles range from saprotrophic to endophytic and obligate biotrophic root-colonizing fungi. Members of Sebacinales are often root-interacting endophytes involved in associations with a very wide range of plant species, and they are globally distributed [2, 3].

*Serendipita indica* colonizes roots of a vast number of different plant species. It is able to enhance the growth and yield of mono- and dicotyledonous plants, and enhances host plant resistance to biotic and abiotic stresses [2, 4–6]. Due to these positive effects on host plants, and its ability to grow in axenic culture, it became a model fungus for studying the physiology and molecular basis of symbiotic plant-microbe interactions [7–10]. It was also suggested for diverse applications, e.g. as a biofertilizer and biocontrol agent [6, 11], for improvement of plant cell culture [12], or *in vitro* cultivated medical plants [13].

*Serendipita indica* can be propagated axenically on various complex and minimal substrates in the absence of host plants [4]. *S. indica* chlamydospores (asexual resting spores) germinate and form hyphae which resemble a string of beads (monilioid hyphae), which, after a few days, develop chlamydospores again.

A range of media have been proposed for axenic *S. indica* propagation, but these media are complex and can be somewhat laborsome to prepare with particular technical requirements [14, 15] or are easy to prepare and economical, but less productive [16]. The most commonly used medium is a Hill-Käfer complex medium (CM) containing 25 different macro-and micro-ingredients [14, 17]. Using this CM agar medium, *S. indica* typically requires 3 weeks to cover a 9.5 cm petri dish with hyphae after plug inoculation [18]. Different modifications of CM medium have been suggested for improving growth of *S. indica* axenically: reduction of the working volume of liquid medium, using different carbohydrate sources and adjustments of their concentrations; and optimizing agitation speed of liquid cultures [15, 19]. However, these efforts added more complexity to medium preparation or growth conditions and had only limited success in increasing fungal growth.

We therefore aimed at developing a growth medium for *S. indica*, which is simple to prepare while providing high inoculum yield. We rationalized that a medium based on plant tissue could be suitable for the facultative saprophyte *S. indica* and started using vegetable juice as a base for the medium. The commercially available V8 juice (Campbell Soup Co.) is the main component of different media used for propagation of plant fungal pathogens, e.g. for sporangia induction in *Phytophthora* [20]. In this study, we describe an optimized medium based on vegetable juice, which is simple to prepare and yields high amounts of inoculum from the root-colonizing model fungus *S. indica*.

## Results

### *Serendipita indica* biomass production in liquid medium

To optimize yield of the plant root colonizing fungus *S. indica*, we tested different media for fungal inoculum production. 10 d after incolation of 100 ml of medium with *S. indica*, the commonly used complex Kaefer-Hill medium (CM) yielded 1.6 g (fresh weight). In comparison, a vegetable juice (VJ) medium enhanced *S. indica* biomass production by a factor of around 13 (Fig. 1a). The basal VJ medium has a simple composition of 15% (v/v) V8-juice (Campbell Soup Corp.; a commercially available vegetable juice) and 5 mM CaCO_3_. Omission of CaCO_3_, which is not only important as a medium component per se, but also to increase the pH to mitigate V8 acidity, led to a significant drop in fungal biomass (Fig. 1a). Similarly, decreasing V8 juice content in the medium from 15% to 7% (v/v) reduced fungal biomass yield significantly (Fig. 1a). A vegetable juice content higher than 15% is not recommended because this leads to a higher viscosity and particle content, which would necessitate additional steps like centrifugation or filtration to retrieve fungal inoculum. Addition of glucose to the VJ medium did not result in a further increase of fungal biomass in liquid culture (Fig. 1a). We also tested a different commercial brand of vegetable juice, which gave comparable results to V8 juice with respect to fungal biomass production (data not shown).

**Figure 1:**
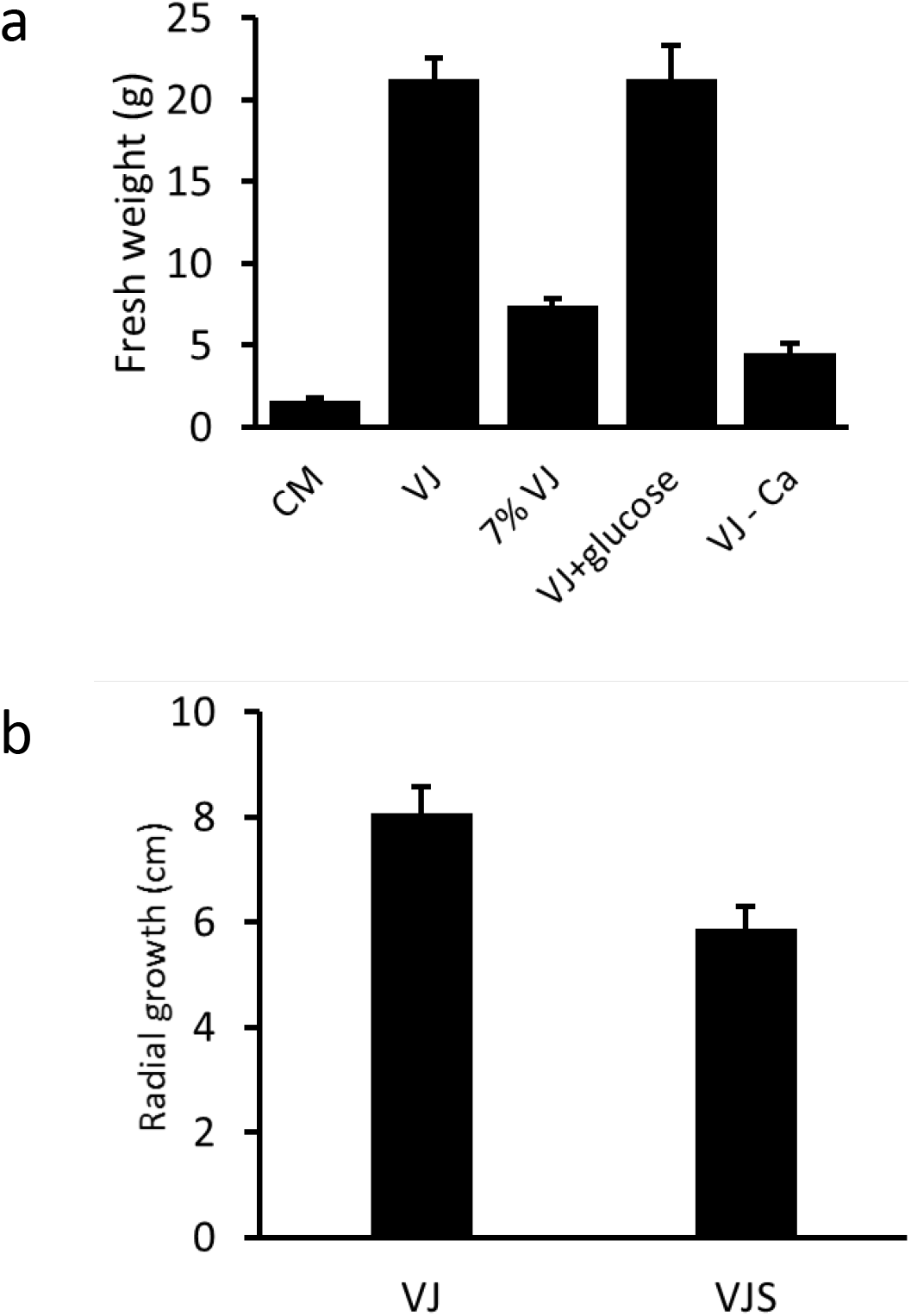
*Serendipita indica* biomass production in different media. a) The fungus was grown in 100 mL of medium in 300 mL Erlenmeyer flasks for 10 days at room temperature (20 ±3 °C) on a shaker (70 rpm). Mycelium was retrieved using Miracloth and fresh weight was determined subsequently. Media used: Complex medium ‘CM’; Vegetable juice ‘VJ’ medium: 150 ml V8 juice, 850 ml distilled water and 5 mM CaCo3; ‘7% VJ’ medium’: VJ medium with 70 ml instead of 150 ml V8 juice; ‘VJ+ D-glucose’: VJ medium supplemented with 10 g/L D-glucose; ‘VJ-Ca’ medium: VJ medium without CaCo3. Values are means of three cultures, with error bars indicating standard deviation. Experiments were independently reproduced at least three times with similar results. b) Diameter of fungal colonies on agar plates prepared with VJ medium or VJS medium. VJS medium is VJ medium supplemented with 40 g/L sucrose and with CaCo3 to a final concentration of 20 mM. Values are means of three cultures, with error bars indicating standard deviation. Experiments were independently reproduced at least three times with similar results.

### Growth of *S. indica* on agar medium for chlamydospore production

To produce *S. indica* chlamydospores for plant root inoculation, *S. indica* is commonly cultivated on agar plates using CM medium. We compared growth of *S. indica* on CM agar with VJ medium. VJ agar plates were completely covered by fungal mycelium 8-10 days post inoculation, while growth on CM medium was significantly slower (Fig. 2a), requiring about three weeks to completely cover the plate with mycelium. Cultivation under continuous light or in darkness had only minor effects on fungal growth, both with VJ and CM medium (Supplementary Fig. 1). Addition of sucrose to the VJ medium did not increase, but rather decrease colony diameter (Fig. 1b). However, sucrose in the VJ medium was required for a high chlamydospore yield on this medium: At 8 dpi, a low number of spores was recovered from *S. indica* grown on CM at this early time point corresponding to the small colony diameter (Fig 2a). Surprisingly, VJ basal medium yielded only few spores although plates were completely covered with fungal mycelium. Adding 4% sucrose and increasing CaCO_3_ concentration to 20 mM in the VJ medium, however, led to a higher spore production compared to both the basal VJ and CM medium (Fig. 2b). We termed this medium VJS (for vegetable juice sucrose medium).

**Figure 2:**
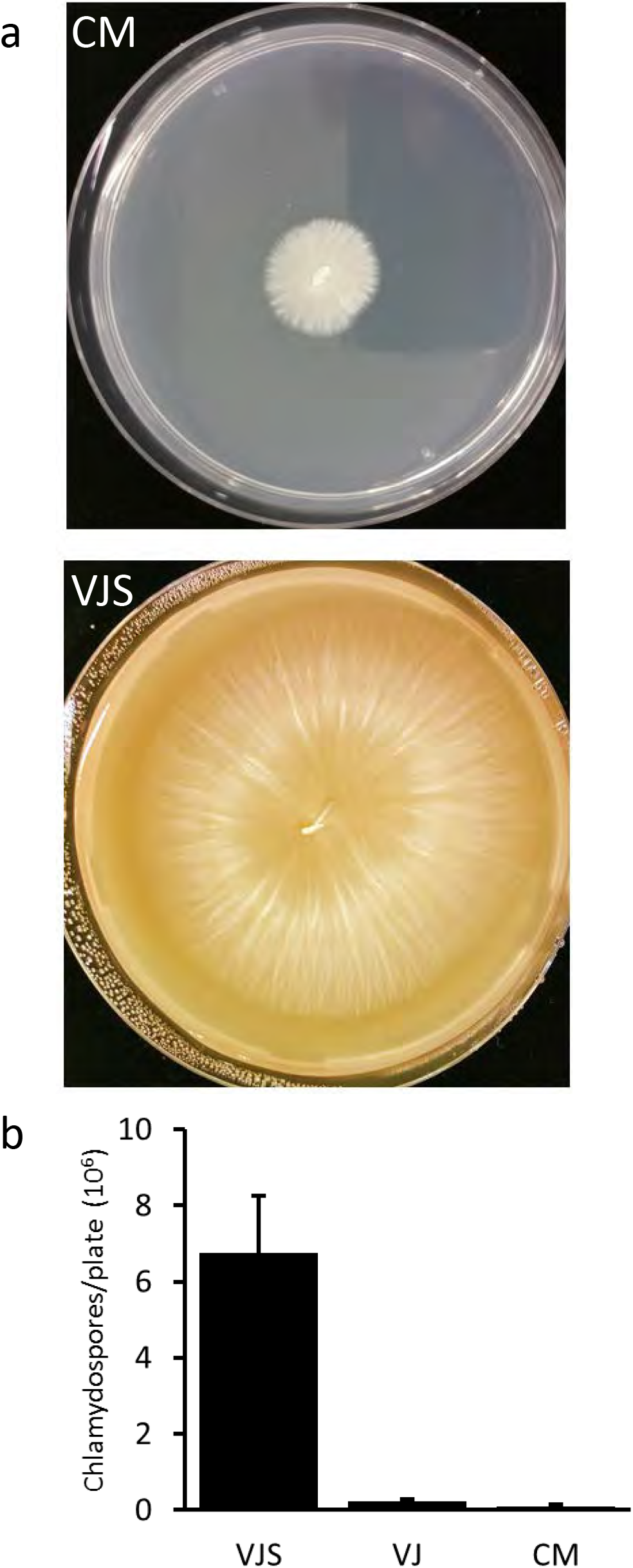
*S. indica* chlamydospore production on agar medium. a) Image of fungal growth on CM agar and VJS agar 8 days after inoculation with an agar plug. Standard petri dishes with a diameter of 9 cm were used and kept at room temperature (20 ±3°C) in the dark. b) Amount of chlamydospores harvested from agar plates. Chlamydospores were obtained by washing the surface of the plates with 15 mL of distilled water using a spatule, and subsequent filtering through Miracloth to remove hyphae and debris. After appropriate dilution, chlamydospores were counted using a microscope and a Fuchs-Rosenthal counting chamber. Values are means from three plates, with error bars indicating standard error. Experiments were independently reproduced at least three times with similar results.

### Effect of Myo-inositol on *S. indica* growth on agar plates

To test the hypothesis that the cause of *S. indica* growth enhancement is because of the Myo-inositol content of V8 [21, 22] a solid medium (yeast extract peptone agar) with 1% Myo-inositol or with 1% glucose was tested. However, the presence of Myo-inositol reduced colony diameter. Moreover, 1% Myo-inositol attenuated aerial mycelia in contrast to the control, which was rich in superficial and aerial mycelia (Supplementary Fig. 2).

### VJ medium is highly suitable for inoculum production of *Fusarium graminearum* and *Verticillium longisporum*

We tested growth of the important wheat pathogen *F.graminearum* on VJ agar medium and recorded the production of macroconidia. A 10-fold increase of macroconidia production was noticed when *F.graminearum* was grown on VJ compared to mung-bean medium (Fig 3). A test with *Verticillium longisporum* revealed that VJ medium was also suitable for production of *V. longisporum* conidia, with one agar plate yielding more than 600 million spores on average.

**Figure 3:**
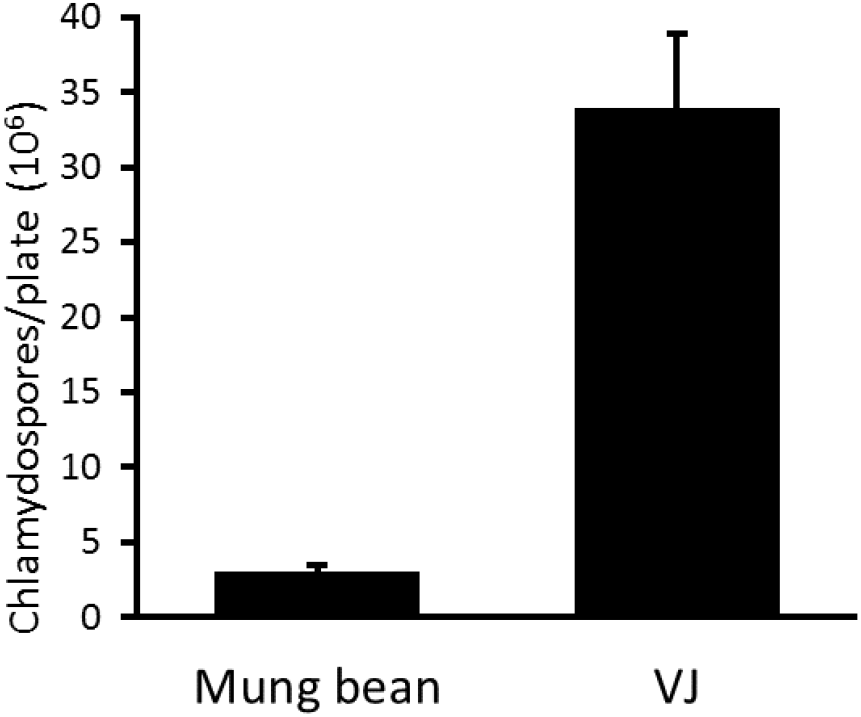
*Fusarium graminearum* macroconidia production on VJ agar. *F. graminearum* was grown on Mung bean agar medium or on VJ agar medium for 14 days in the dark. Total number of macroconidia recovered from a single plate is presented. Values are means from three plates, with error bars indicating standard error. Experiments were independently reproduced at least three times with similar results.

### Comparison of ultrastructural features between CM- and VJ-grown *S. indica*

To be able to identify possible differences in the ultrastructure of *S. indica* hyphae grown in CM and VJ medium, we prepared hyphae from *S. indica* liquid cultures grown for 11 days for electron microscopy. Hyphae from both media appeared very similar with respect to size and ultrastructural features. Samples from both media showed similar densities of nuclei, and of lipid bodies (Figure 4).

**Figure 4:**
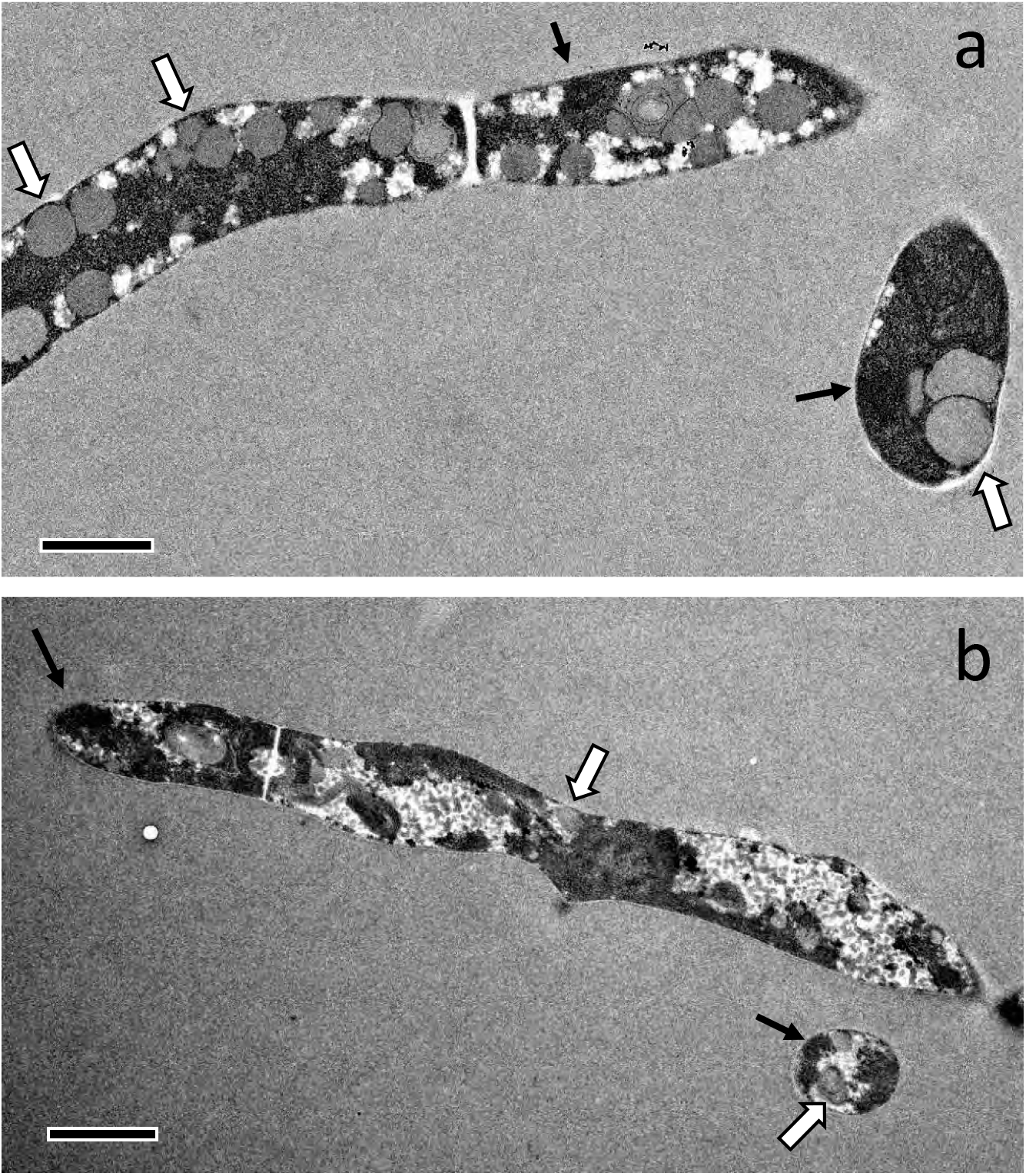
Morphology of *Serendipita indica* hyphae from CM and VJ medium. *S. indica* was grown in 100 mL of CM medium (a) or VJ medium (b) for 10 days at room temperature (20 ±3 °C) on a shaker (70 rpm), and hyphae were subsequently harvested for fixation and preparation of electron microscopic images. Hyphal diameters were between 1 and 2 μM in both media. Black arrows indicate nuclei, white arrows indicate lipid bodies. Bars represent 1 μm (a) and 2 μm (b).

### Interaction of VJS-grown *S. indica* chlamydospores with plants

Using chlamydospores from VJS medium, we inoculated roots of 7-day-old barley plants. Staining of these roots 7 days after inoculation using WGA-Alexafluor 488, we observed fungal hyphae as well as chlamydospores colonizing the roots (Supplementary Fig. 3a), similar to published reports on barley root colonisation [5, 8]. We also used *S. indica* plugs from VJS medium and inoculated 7 day-old Arabidopsis thaliana (Col-0) seedlings growing on ½ MS medium (Supplementary Figure 3b). Presence of the fungus induced growth of the seedlings, visible 7 days after inoculation, comparable to published reports using *S. indica* fungal plugs grown on CM medium [23].

## Discussion

We aimed at developing a medium that is easy to prepare, with only few and readily available, inexpensive ingredients which provides high inoculum yields for the fungus *Serendipita indica*, a fungus used as a model system to study plant-endophyte interactions in the lab and, due to its growth enhancing and plant strengthening properties, also studied in agricultural and biotechnological settings.

We optimised a medium for *S. indica* using commercially available vegetable juice as the main ingredient. Media based on the commercially available vegetable juice V8 have been shown to be useful for growth and sporangia induction of the plant pathogenic fungus *Phytophthora* [20], and are frequently used for conidia production in different ascomycetes including *Fusarium culmorum, Parastagnospora nodorum* and *Pyrenophora tritici-repentis* [24, 25].

In a series of experiments, we determined that a medium composition of 15% (v/v) V8 vegetable juice and 0.1-0.4% (w/v) CaCO_3_ are optimal, and superior to traditionally used complex medium (CM) with respect to the speed of hyphal growth and fungal biomass production in liquid culture. The observed high growth rate of *S. indica* did not depend on the brand of vegetable juice used, as comparable results were obtained when a different brand of vegetable juice was used (with tomato juice as the main component, and carrot, beetroot, celery, fermented cabbage, leek, parsley and lemon juice). Fungal inoculum obtained from liquid culture is frequently used for inoculation of larger plants, e.g. in pot experiments with barley or field experiments with wheat [5, 26] and we propose VJ medium for fast and efficient inoculum production. Pham et al. (2004) reported yields between 3.67 and 2.39 g fresh weight per 100 mL in CM medium, similar to our results shown in Fig. 1a. A report on enhancing *S. indica* biomass yield, using a modified Kaefer medium in a 14-L bioreactor, achieved production rates of 0.18g L^-1^h^-1^ [14]. This is about four times less than our results, using 300 ml Erlenmeyer flasks with 100 ml culture volume, yielding 21 g after 10 d of culture (Fig. 1).

In other experiments, e.g. when a defined amount of sterile fungal inoculum is required, *S. indica* chlamydospore preparations have been used [8, 27, 28]. For this purpose, chlamydospores can be easily retrieved by washing agar plates with water. A high chlamydospore yield was obtained with agar plates using VJ medium supplemented with 4% Sucrose (VJS medium) (Fig. 2). Even higher chlamydospore yields may be obtained by using liquid culture in a bioreactor and subsequent removal of hyphae, as shown by [14], with yields up to 9.25*10^7^ spores per mL. Nevertheless, as fungal cultures on agar plates do not require specific equipment and hardly any hands-on time, it will probably remain the method of choice, at least for chlamydospore amounts required for research purposes.

We confirmed that *S. indica* inoculum produced with VJ medium has similar ultrastructural features as CM medium-grown fungus (Fig. 4). Roots were well colonised by the fungus after inoculation with VJS-grown fungus (Supplementary Fig. 3a), and *S. indica*-induced growth promotion was observed for Arabidopsis plants (Supplementary Fig. 3b), in line with previous reports on the properties of the fungal endophyte when propagated on CM medium [23, 29]. We therefore suggest VJS medium for efficient chlamydospore production. In addition, VJ medium is also highly suitable for inoculum production of *Fusarium graminearum* and *Verticillium longisporum:* Macroconidia production of *F. graminearum* was about 10 times higher than with traditionally used mung bean agar (Fig. 3).

Some studies explained high growth rates on V8-based media with the presence of an inducing factor in V8 juice. However, this inducing factor was later found to be rather a combination of factors required for induction of sexual development of *C. neoformans* [30]. We speculate that beneficial properties of VJ medium for a high growth rate of *S. indica* are due to its complex nutrient composition and possibly the presence of secondary plant metabolites present in vegetable juice.

## Conclusions

The optimized VJ medium described here was superior to complex media commonly used for production of fungal inoculum of the plant root endophytic model fungus *Serendipita indica*. Due to its ease of preparation and faster and more efficient inoculum production as compared with Kaefer-Hill medium, we expect this medium to become the medium of choice for *S. indica*. We further show that other fungal inocula used in plant pathology can also be produced efficiently using VJ medium. Besides its usefulness for research purposes due to the accessibility and relatively low price of its components, and the ease of preparation, it may also serve as a basis for the development of large-scale inoculum production for commercial purposes.

## Methods

### Fungal materials

*Serendipita indica / Piriformospora indica* isolate DSM11827 was obtained from the DSMZ (Leibniz-Institut DSMZ-Deutsche Sammlung von Mikroorganismen und Zellkulturen GmbH, Braunschweig, Germany). Fusarium strain *F. graminearum* was provided by Prof. Dr. R. Hückelhoven, Technical University Munich. *Verticillium longisporum* strain Vl43 [31] was obtained from Prof. W. Dröge-Laser, Würzburg University.

### Media preparation

VJ medium was prepared using the commercially available vegetable juice V8 (‘V8 Original Vegetable Juice’; Campbell Soup Company, Camden NJ, USA), which is mainly composed of tomato juice (86%), supplemented with juice from beetroot, carrot, celery, spinach, parsley, lettuce and watercress, NaCl (6.8 g/L), and natural flavouring (www.v8juice.co.uk). For one liter of VJ medium, 850 ml deionized, distilled water was supplemented with 150 ml of V8 vegetable juice and 0.5 g CaCO_3_. For VJ medium supplemented with sucrose (VJS medium), we used 20 mM of CaCo3 and added 40 g/L sucrose to the VJ medium.

For *S. indica* liquid culture, 300 ml Erlenmeyer flasks containing 100 ml of the VJ medium were used. The flasks were inoculated with 2 plugs (5 mm diameter) from agar plates containing *S. indica* and kept at 20 °C on a rotary shaker (70 rpm) for 10 days and harvested by filtration through miracloth. For production of *S. indica* chlamydospores on solid medium, VJS medium was supplemented with 1.2% (w/v) Phyto agar (Duchefa-biochemie.com). CM medium was based on the Aspergillus medium [32], as described in [15] without casamino acids and without agar for liquid medium, while 1.2%Phyto agar was used for solid medium in petri dishes. Mung bean agar was prepared as described in [33] with 1.2%Phyto agar. Arabidopsis seedlings were grown on ½ MS medium (Murashige and Skoog medium basal mixture, including MES buffer, pH 5.8; Duchefa Biochemie B.V.) containing 0.5% sugar and 1 %Phyto agar. To test the impact of myo-inositol, one liter of medium was prepared with distilled water, tryptone (10 g), yeast extract (5 g) and micro elements (1 ml micro-element solution of the Aspergillus medium as described in [15]), supplemented with 10 g D-glucose or 10 g myo-inositol.

### Electron microscopy

Spores and hyphae of *S. indica* grown in liquid medium for 11 days were centrifuged and the supernatant was removed. The resulting fungal material was fixated with 2.5%glutaraldehyde in cacodylate buffer over night at 4 °C. Further steps of electron microscopy sample preparation were as described in [34]. Images were recorded on a JEM-2100 transmission electron microscope (JEOL, Freising, Germany) at 200 kV acceleration voltage with a F416 digital camera TVIPS TemCam F416 (Tietz Video and Images Processing Systems, Gauting, Germany).

### Fluorescence microscopy

For microscopic investigation of fungal root colonization, barley roots were stained with Wheat Germ Agglutinin-Alexafluor 488 (WGA488; Molecular Probes; Invitrogen, www.invitrogen.com) as described in Deshmukh et al. (2006). Images were recorded on a confocal laser scanning microscope (Leica TCS SP5) using the bright field channel and a GFP filter set for detection WGA488.

## Authors’ contributions

MO and FW designed the study. MO, CS and FW performed experiments, all authors analyzed and interpreted the data. The manuscript was prepared by MO and FW, and all authors contributed to and approved the final version of the manuscript.

## Acknowledgements

We thank Paula Puente for preparing *S. indica*-colonised barley plants. Fusarium strain *F. graminearum* was kindly provided by Prof. Dr. R. Hückelhoven, Technical University Munich. *Verticillium longisporum* was kindly provided by Prof. W. Dröge-Laser, Würzburg University.

