## Supplementary Figures 1-3 for "An improved growth medium for enhanced inoculum production of the plant growth-promoting fungus *Serendipita indica*"

### Supplementary Figure 1

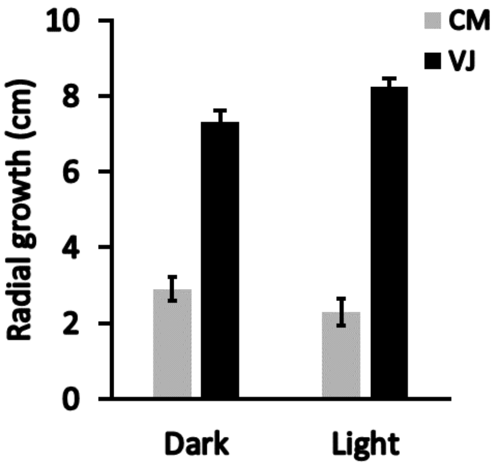


**Supplementary Figure 1: Diameter of *S. indica* hyphal growth on CM and VJ agar**

Diameters were determined 8 days after inoculation of CM or VJ plates with an *S. indica* agar plug. Standard petri dishes with a diameter of 9 cm were used and kept at room temperature (20 ±3 °C) either in the dark or in continuous light. Values are means from three plates, with error bars indicating standard deviation. Experiments were independently reproduced two times with similar results.

### Supplementary Figure 2

a

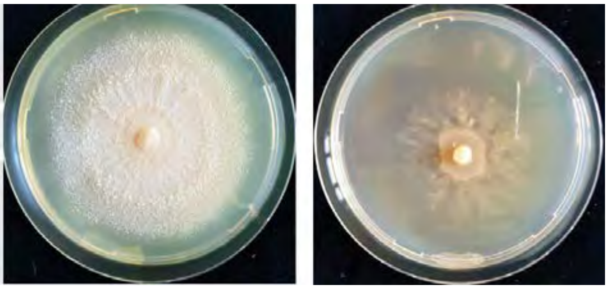

b

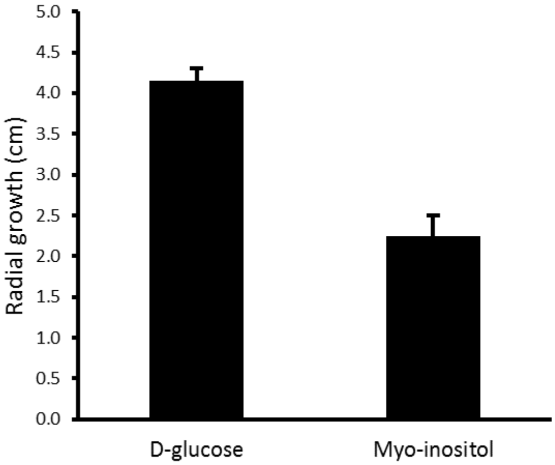


**Supplementary Figure 2: Effect of Myo-inositol and Glucose on *S. indica* growth**

a) Image of fungal growth on tryptone yeast agar supplemented with 10 g/ L D-glucose (left) or 10 g/L myo-inositol (right) 8 days after inoculation with an *S. indica* agar plug. Standard petri dishes with a diameter of 9 cm were used and kept at room temperature (20 ±3 °C) in the dark.

b) Fungal growth on agar plates supplied with 10 g/L D-glucose or 10 g/L myo-inositol 8 days after inoculation. Values are means from three plates, with error bars indicating standard deviation. Experiments were independently reproduced at least two times with similar results.

### Supplementary Figure 3

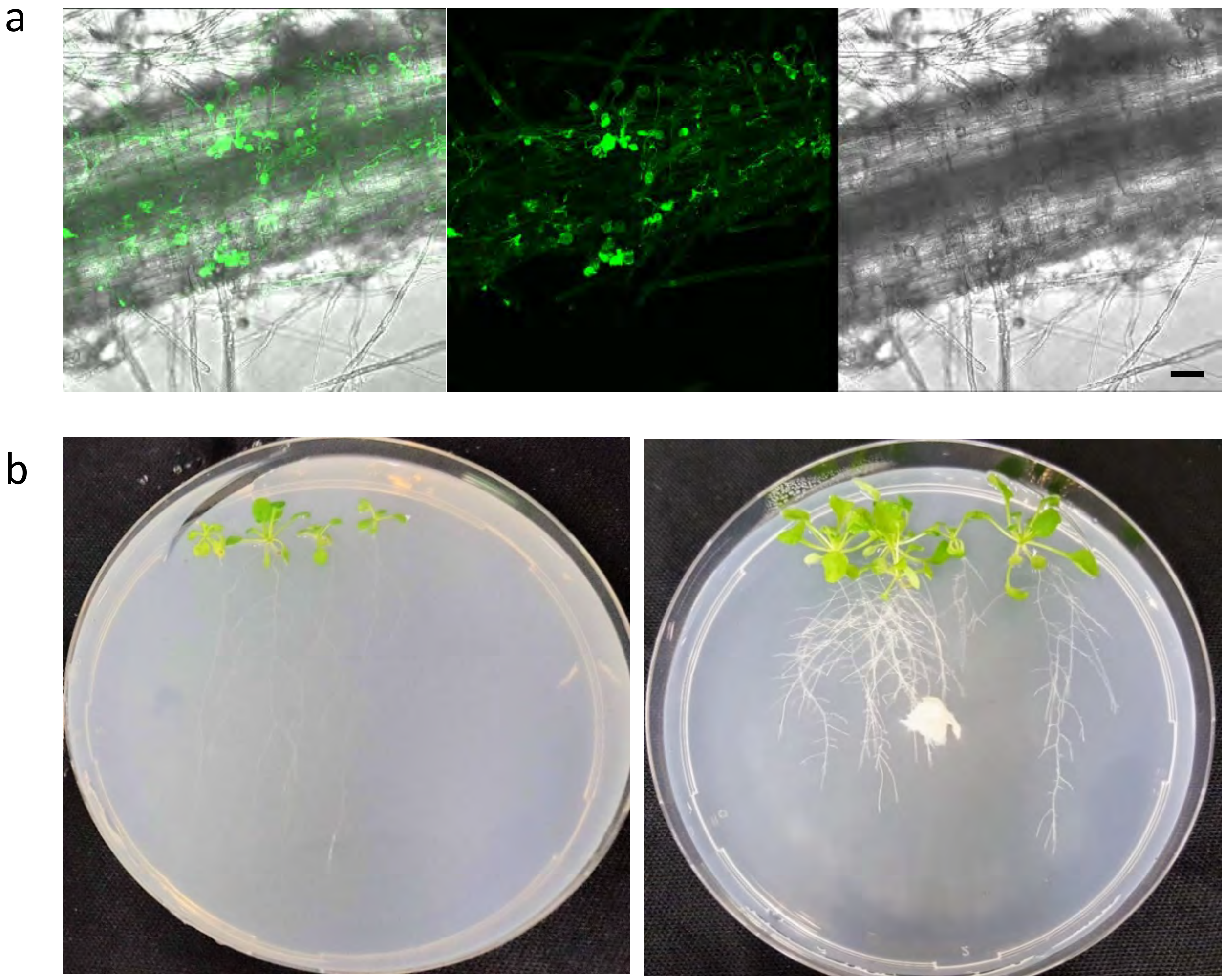


**Supplementary Figure 3: Plant interaction of *S. indica* from VJ-grown inoculum**

a) Colonisation of barley roots with *S. indica* after inoculation with spores prepared from VJS medium. Roots of 7 d-old seedlings grown in hydroponic medium were inoculated and analyzed 7 d after inoculation. Germinated chlamydospores with their hyphae are visualized by staining root sections with a fluorescent dye-coupled wheat germ agglutinin (WGA-AF 488). Images were recorded using a confocal fluorescence microscope. Shown are the bright field image (right), fluorescence image (middle) and an overlay of these two images (left). The size bar represents 50 µm.

b) Growth of Arabidopsis thaliana (Col-0) plants in the absence (left) or presence (right) of *S. indica*. One-week-old Arabidopsis seedlings growing in petri dishes (9 cm diameter) on ½ MS medium were inoculated with a *S. indica* mycelial plug obtained from VJS medium. Plates were photographed after 3 weeks of growth under short day conditions (8 h light / 16 h dark, 22°C).
